# Computational modeling of the photon transport, tissue heating, and cytochrome C oxidase absorption during transcranial near-infrared stimulation

**DOI:** 10.1101/708362

**Authors:** Mahasweta Bhattacharya, Anirban Dutta

## Abstract

Transcranial near-infrared stimulation (tNIRS) has been proposed as a tool to modulate cortical excitability. However, the underlying mechanisms are not clear where the heating effects on the brain tissue needs investigation due to increased near-infrared (NIR) absorption by water and fat. Moreover, the risk of localized heating of tissues (including the skin) during optical stimulation of the brain tissue is a concern. The challenge in estimating localized tissue heating is due to the light interaction with the tissues’ constituents, which is dependent on the combination ratio of the scattering and absorption properties of the constituent. Here, apart from tissue heating that can modulate the cortical excitability (“photothermal effects”); the other mechanism reported in the literature is the stimulation of the mitochondria in the cells which are active in the adenosine triphosphate (ATP) synthesis. In the mitochondrial respiratory chain, the complex IV, also named as the cytochrome c oxidase(CCO), is the unit four with three copper atoms. The absorption peaks of CCO are in the visible (420-450nm and 600-700nm) and the near-infrared (760-980nm) spectral region which have been shown to be promising for low level light therapy (LLLT), also known as “photobiomodulation”. While much higher CCO absorption peaks in the visible spectrum can be used for the photobiomodulation of the skin, 810nm has been proposed for the non-invasive brain stimulation (using tNIRS) due to the optical window in the NIR spectral region. In this article, we applied a computational approach to delineate the “photothermal effects” from the “photobiomodulation,” i.e., to estimate the amount of light absorbed individually by each chromophore in the brain tissue (with constant scattering) and the related tissue heating. Photon migration simulations were performed for motor cortex tNIRS based on a prior work that used a 500mW cm^−2^ light source placed on the scalp. We simulated photon migration at 630nm and 700nm (red spectral region) and 810nm (near-infrared spectral region). We found a temperature increase in the scalp below 0.25 ° C and a minimal temperature increase in the gray matter less than 0.04 ° C at 810nm. Similar heating was found for 630nm and 700nm used for LLLT, so photothermal effects are postulated to be unlikely in the brain tissue.

## 1. Introduction

Near-infrared (NIR) light has been reported to be able to penetrate the extra-cranial layers like scalp, skull, cerebrospinal fluid and reach the superficial layers of the cerebral cortex due to the optical window. It has been hypothesized that interaction of NIR light with cytochrome c oxidase (CCO) can potentiate the CCO in the mitochondria, a component of the electron transport chain and key complex in energy production[1]. CCO is the primary chromophore in the mitochondria besides the calcium-ion channel (possibly mediated by opsin light absorption). Secondary effects of the photon absorption include ATP increase, brief explosion of reactive oxygen species, an increase in nitric oxide, and calcium levels modulation. Tertiary effects include activating a wide range of transcription factors that lead to improved cell survival, increased proliferation and migration, and synthesis of proteins. The interaction of photons with CCO has been found primarily due to the photoacceptor of the binuclear copper center(CuA)[2] in the NIR (700-980nm) range[3]. CCO accepts photons and transduces photo-signal in the NIR spectrum[4] which is postulated to be the underlying mechanism of "photobiomodulation" (PBM). The underlying theory suggests that nitric oxide (NO), which inhibits the enzymatic activity of CCO, can be dissociated by photons absorbed by the CCO that has two heme and three coppers with different absorption spectra[5]. The dissociation of the inhibitory NO[5], thereby allowing respiration to resume unhindered, increasing energy production (ATP synthesis) [6]. Consequently, various signaling molecules are activated, including (but not limited to) ROS, cyclic adenosine monophosphate(cAMP), NO, and calcium. While the underlying mechanisms are still elusive, it has been seen that the increase in reactive oxygen species (ROS) during PBM may have the ability to trigger mitochondrial signaling pathways which leads to cytoprotective, anti-oxidant and anti-apoptotic effects in cells[7].

Furthermore, one can not only increase the activity of CCO (primarily at 810nm) but can also reduce the activity of isolated CCO using two NIR wavelengths (750nm and 950nm)[8]. Therefore, the light wavelength is important since the efficiency of red (600-700nm) to NIR (700-980nm) spectrum varies due to their varied ability to modulate CCO and the energy production [9]. The Cu^2+^ centers of CCO are assumed to be one of the causes of the CCO interaction with red and NIR light[10]. However, CCO shows much higher absorption around the 420 and 450nm[11] in the visible range due to the two heme groups a and a3[12]. Blue and green light have shown promises in stem cell differentiation[13] where the effect can be due to light-sensitive ion-channels besides PBM since the transition from glycolysis to oxidative phosphorylation is also a crucial factor in stem cell differentiation. Nevertheless, visible spectrum, especially in the blue and green range, poses a considerable challenge in its utility for targeting deeper tissues due to their low penetration depth [14] where red-NIR spectral range performs better for non-invasive brain stimulation due to the optical window [15]. In the red-NIR spectral range, the red spectrum has a lower penetration depth; hence, it is more efficient for skin[16] or other surface tissues, whereas tNIRS is better suited for non-invasive brain stimulation [1].

In this article, we investigated the red-NIR spectral region with the CCO absorption peaks selected in the range of 600-700nm for red and 760-980nm for NIR. Here, an average power density of 5mW cm^−2^ to 500mWcm^−2^ on the surface of the skin is used for non-invasive brain stimulation. However, there is a pronounced biphasic dose response, with low light levels having stimulating effects, while high light levels have inhibitory effects[14] that needs biochemical investigation in conjunction with computational modeling. As the power density increases, the photothermal effects needs consideration besides photobiomodulation due to increased tissue heating, which can affect the biochemical responses and brain excitability. Here, the optical energy leading to tissue heating is based on the fundamentals of increased absorption of longer wavelengths by water in the tissue[17][18]. Indeed, photothermal neurostimulation using the 1064nm laser at the frontal cortex has been shown to improve cognitive functions along with neurometabolic activity[9]. Moreover, photothermal neurostimulation has been shown to be promising to map mesoscale brain connectomes[19].

A significant advantage of PBM over photothermal stimulation for non-invasive brain stimulation is that it is safe without the heating effects and can be advantageous therapeutically[3]. Here, tNIRS has been shown to increase cerebral blood flow, greater oxygen availability, higher oxygen consumption, improved ATP production, and enhanced mitochondrial activity. PBM has been found to be safe and well-tolerated as a potential treatment of depression, anxiety, and cognitive impairment[20][21][22]. Cognitive ability has also been shown to improve after several months of treatment by the light emitting diode at 633nm (red) and 870nm (near-infrared) in patients with chronic traumatic brain injury[23]. Also, tNIRS has been reported as a possible treatment for ischemic stroke with an application at 808nm (near-infrared laser)[24]. Moreover, the use of transcranial NIR laser (810nm) in low level light therapy showed improvement in patients suffering from anxiety and depression[25]. Since increased oxygen consumption occurs during increased neural activity[26], which leads to increased CCO activity, so an assessment and modulation of CCO activity can open a pathway to monitor and modulate neuronal activity[27] [28]. Here, redox state-dependent changes in the NIR spectrum is an essential tool for near-infrared spectroscopy of the oxidation state of CCO[2].

In this paper, investigation of the interaction of light with the chromophores that are responsive to photons in the red-NIR spectral region has been performed. Since there is an increased absorption of longer wavelengths by water in the tissue, we postulate that NIR light interaction with neural tissue may have effects of photobiomodulation as well as photothermal neurostimulation which needs consideration for rational dosing of tNIRS due to the biphasic dose response. Although it has been reported that tNIRS is a modulator of cortical excitability in healthy human brain[1] which forms the basis of this paper, however, the exact mechanisms of the neuromodulation has been elusive. In this paper, we apply computational modeling to dissociate photothermal effects from photobiomodulation during tNIRS with 810nm while comparing that with the red spectrum (630nm and 700nm) used for LLLT[16] to investigate the mechanisms underlying neuromodulation [1]. Here, the primary aim is to better understand the extent of optically induced tissue heating (primarily due to water and fat absorption) during tNIRS based on the experimental results by Chaieb et al. [1].

## 2. Methods

### 2.1. Head Model Selection

To develop the computational model of light interaction with the chromophores in the human head by non-invasive approach, a digital brain phantom based on high-resolution brain atlas[29] was used in the study. From the Colin27 head atlas, the different layers of the brain were segmented to form layered tissues of the head model[30]. Volume mesh was created from each layer after surface smoothing using CGAL surface mesh toolbox[31] with each surface having its own mesh criteria and density. After the multi-layered surface mesh was generated, the volume mesh was generated using the Delaunay tetrahedralization algorithm[32]. Figure 1 shows the multi-layered head mesh generated after Delaunay tetrahedralization.

**Figure 1.**
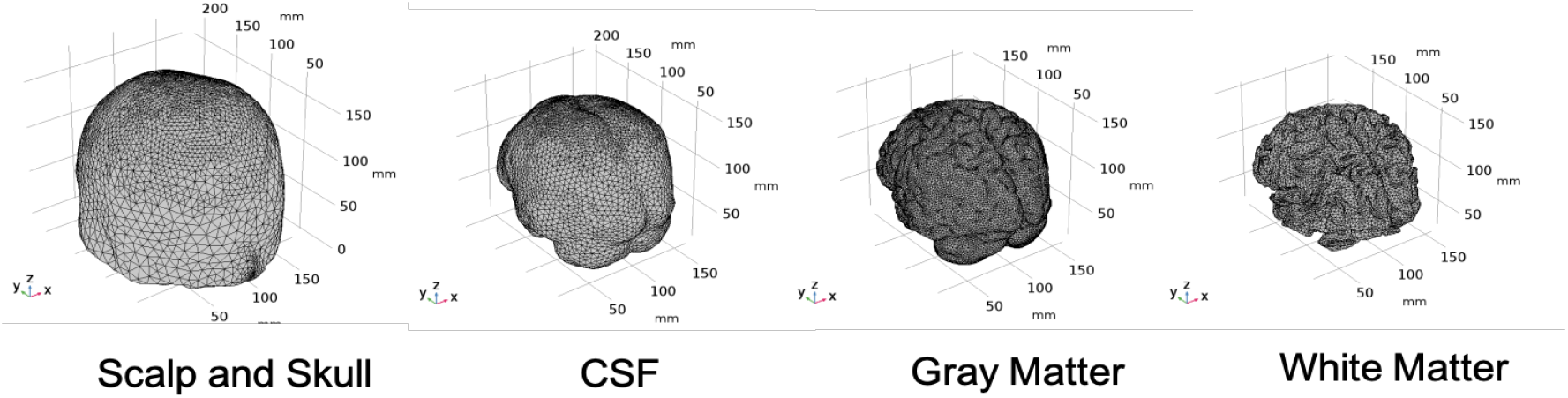
Tetrahedral Mesh for the four-layered Colin27 Head Model

The details of the mesh components of the complete four-layered Colin27 head model is given in table 1.

**Table 1.**
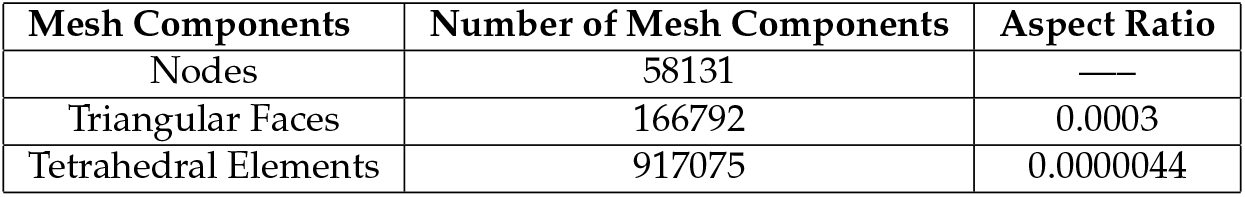
Details of Mesh Components of the four-layered Colin27 Head Model

| Mesh Components | Number of Mesh Components | Aspect Ratio |
| --- | --- | --- |
| Nodes | 58131 | — |
| Triangular Faces | 166792 | 0.0003 |
| Tetrahedral Elements | 917075 | 0.0000044 |

### 2.2. Geometry and domain assignment

The tetrahedral mesh was imported to COMSOL, a Finite Element Method(FEM) simulation software, as a CAD model and each layer was designated as a domain and corresponding optical properties were assigned to each domain. The domains are as follows:

- Domain 1: Combined scalp and skull
- Domain 2: Cerebrospinal Fluid
- Domain 3: Gray Matter
- Domain 4: White Matter

### 2.3. Simulation of Radiative Transfer Equation using Diffusion Approximation

Photon transport in scattering media, such as biological tissues, is generally modeled using the radiative transfer equation (RTE)[33] due to its more accurate solution for highly scattering medium as in the case of brain tissues [34] and higher computational efficiency for complex medium[35][36]. The tetrahedral mesh generated was converted into a CAD file and imported to COMSOL. After importing the mesh to COMSOL, the computation of photon propagation was solved through the diffusion approximation of the RTE. The second order partial differential equation(eqn. 1) describes the time behavior of photon fluence rate distribution in a low-absorption high-scattering medium.

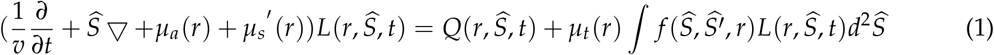

Here, *µ*_*a*_, 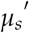, and *µ*_*t*_ are the absorption, reduced scattering and total attenuation coefficients, respectively, 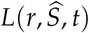, the radiance at position r with direction of propagation S, v the velocity of light through the medium (v = c/n where c is the velocity of light in vacuum and n the refractive index of the medium), 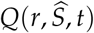 the source term, and 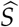, 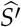, *r* the phase function for scattering. A standard approximation method for the RTE assumes that the radiance in tissue can be represented by an isotropic fluence rate, *φ*(r,t), plus a small directional flux, J(r,t), where:

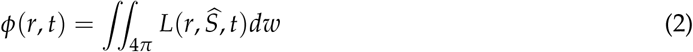

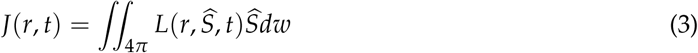

The final diffusion approximation of RTE, i.e., diffusion equation, is derived as:

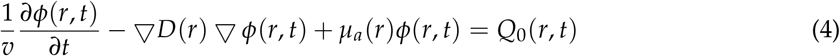

where D(r) is defined as:

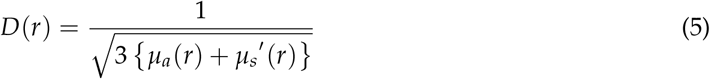

The 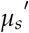 is the reduced scattering coefficient and is obtained from equation 6:

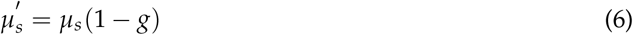

where *g* is the anisotropy factor.

The diffusion equation is solved by using the COMSOL Multiphysics software using the Partial Differentiation Equation (PDE) toolbox (comparison with Monte Carlo simulation shown in the supplementary material 1). The entire head model had four domains: scalp and skull combined, CSF, gray matter, and white matter, as listed in table 2. The physics was applied to each domain at steady state with the initial condition being zero. The source term was taken from the published literature[1] where the power density was 500mW/cm^2^ at the scalp surface as presented by Chaieb and colleagues[1]. The head model was assumed to be surrounded by air at room temperature(25°C). We placed our sources at the air-tissue interface, which is at the scalp, following Chaieb and colleagues[1]. The boundary condition here is as follows:

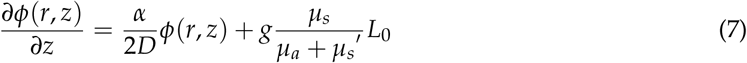

**Table 2.** Whole tissue optical properties of each layer in the head model at the three wavelengths

| Domains | Absorption Coefficient<br>(1/m) |  |  | Reduced Scattering Coefficient<br>(1/m) |  |  |
| --- | --- | --- | --- | --- | --- | --- |
|  | 630nm | 700nm | 810nm | 630nm | 700nm | 810nm |
| Scalp and Skull | 19 | 13 | 16 | 858 | 900 | 760 |
| Cerebrospinal Fluid | 4 | 4 | 2.6 | 250 | 250 | 250 |
| Gray Matter | 127.25 | 62.91 | 57.09 | 990 | 880 | 746 |
| White Matter | 66.11 | 32.52 | 20.77 | 4400 | 4356 | 4070 |

The optical properties, namely scattering coefficient at these wavelengths have been reported in various prior works [37] [38]. The absorption coefficients of the tissues are calculated as the summation of the absorption coefficient due to the contribution of each component of interest in the corresponding tissue[39]. The optical properties for the whole tissues at the three wavelengths used in the study are given in table 2.

The reduced scattering coefficients are calculated based on the scattering coefficient and the anisotropy factor (eqn. 6). The anisotropy factor, *g* = 0.89 has been assumed for all the tissue layers. Although literatures have shown that diffuse reflection occurs at the skin surface, in this paper, the reflection effects have been excluded.

### 2.4. Optically induced thermal effects modeled using bio-heat transfer mechanism

The thermal effect due to the absorbed incident light is modeled using the bio-heat transfer mechanism[40]. The algorithm analyzes the temperature distribution and heating profile when the heat is applied to the tissue. The Pennes Bio-heat equation(eqn. 8) is used to model this phenomenon for localized and distributed energy source.

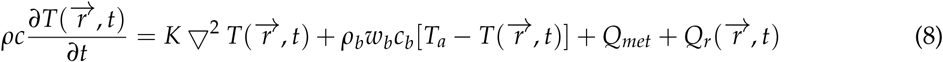

Here, *ρ*(kgm^*−*3^) is the tissue density, c is specific heat of the tissue(kJ/kg/K), K is thermal conductivity, cb(3664J/kg.°C) is blood specific heat, *ω*_*b*_ is blood volumetric perfusion rate, *T*_*a*_ is the arterial blood temperature(37°C), *ρ*_*b*_(1050kg*m^−^*^3^) is the blood density and *Q*_*met*_ and *Q*_*r*_ are the volumetric metabolic heat and the external spatial heating respectively. The heat source term is related to the local fluence rate and tissue absorption coefficient[41] as follows:

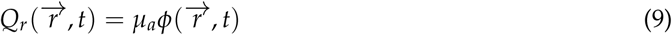

The bio-heat physics is applied with the different tissue components having their respective thermal and blood perfusion properties. The thermal and the blood perfusion parameters are taken from[42] to be used for the computation of the bio-heat transfer, as shown in table 3 and 4.

**Table 3.**
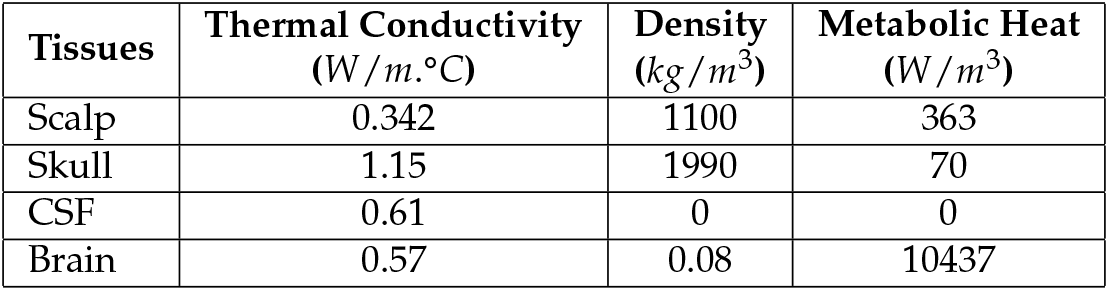
Thermal Properties of Brain Tissues

**Table 4.** Blood perfusion parameters for layers in the head model

| Tissues | Blood Specific Heat<br>($J/kg.K$) | Blood Perfusion<br>($1/s$) | Blood Density<br>($kg/m^3$) | Metabolic Heat Source<br>($W/m^3$) |
| --- | --- | --- | --- | --- |
| Scalp | 3600 | 0.00143 | 1050 | 363 |
| Skull | 3600 | 0.000143 | 1050 | 70 |
| Cerebrospinal Fluid | 3600 | 0 | 1050 | 0 |
| Brain | 3600 | 0.08 | 1050 | 10437 |

The boundary condition was applied at the skin surface. It was assumed that there is heat loss at the skin surface by convection to ambient[42]. For the whole scalp(skin), the convective heat flux value was assumed to be 4*W*/*m*^2^.°*C*[43].

### 2.5. Investigation of Individual Chromophore Absorption in the Tissue

Absorption of red or near-infrared photons by cytochrome C oxidase (unit IV of the mitochondrial respiratory chain) has been established by prior works[44][45]. In this section, we investigated the absorption by CCO at 630nm, 700nm, and 810nm wavelengths, which can cause its activation and may lead to photobiomodulation in the gray matter[44]. Besides CCO, we also investigated other major chromophores in the gray matter along with the investigation of water absorption. Chromophores present in the gray matter that were investigated in this section are as follows:

- Oxyhemoglobin
- Deoxyhemoglobin
- Cytochrome c oxidase(reduced and oxidized state)
- Lipid

Figures 2[46],3[47],4[48] and 5[49] shows the absorption spectra of two states of hemoglobin, two states of cytochrome c oxidase and lipid respectively. In figure 4, the main plot shows the absorbance due to 4.9*µ*M of CCO and the inset shows the absorbance due to five times the concentration of CCO[48].

**Figure 2.**
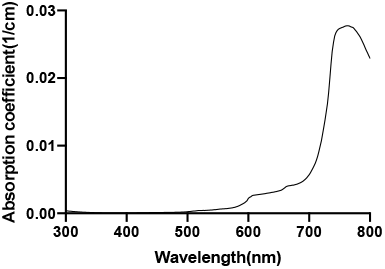
Water absorption spectrum

**Figure 3.**
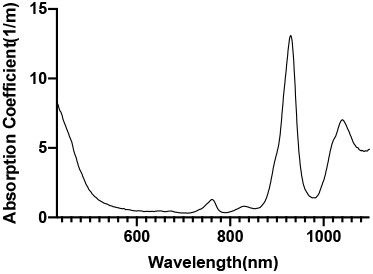
Lipid absorption spectrum

**Figure 4.**
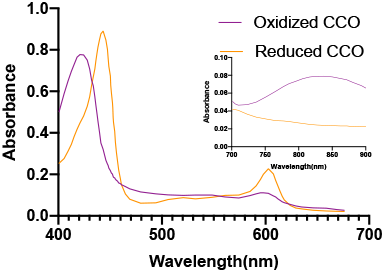
Cytochrome c Oxidase absorption spectrum

**Figure 5.**
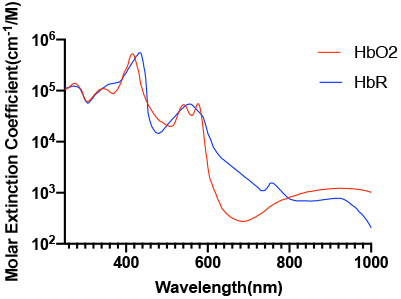
Hemoglobin absorption spectrum

### 2.6. Optical Parameters of Individual Chromophores

The three wavelengths, 630nm, 700nm, and 810nm in the red and the NIR spectral regions, have been reported to be promising for photobiomodulation[44]. The two reported wavelengths were chosen from red (630nm, 700nm) and one in NIR (810nm) spectral region.

The water has very low absorption in the red and near-infrared spectrum, although it increases with increasing wavelength. Calculation of tissue absorption specifically due to water was performed by obtaining the value of the absorption coefficient of pure water at the three wavelengths[46][50][51]. Since 75% water per unit volume (i.e., volume fraction) is present in brain tissues, hence, 0.75*mu*_*a*_ is the absorption coefficient of the tissue specifically due to water[52] [more details provided in the supplementary materials]. The percentage of the dry weight of lipid in gray and white matter[53] (i.e., mass fraction), density of gray and white matter along with the specific absorbance of lipid[47] contribute to the absorption coefficient of the tissue specifically due to lipid. The absorption due to hemoglobin in the brain tissue is dependent on cerebral blood volume[54] and was calculated based on the volume fraction of the blood in the cerebral tissue, hemoglobin oxygen saturation of mixed arterio-venous vasculature, and the absorption coefficient of pure oxy and deoxyhemoglobin [55]. The molar concentration of oxidized and reduced CCO (in mM) were first obtained for the gray and white matter[56]. The absorption coefficient of 1mM of CCO was obtained [2] based on which the tissue absorption coefficient due to oxidized and reduced CCO was calculated. The calculations of the absorption coefficients are provided in the Supplementary Materials.

**Table 5.** Absorption Coefficient of gray and white matter due to the specific chromophores based on their concentration

| Components | Absorption Coefficient<br>(1/m) |  |  |  |  |  |
| --- | --- | --- | --- | --- | --- | --- |
|  | Gray Matter |  |  | White Matter |  |  |
|  | 630nm | 700nm | 810nm | 630nm | 700nm | 810nm |
| Water | 0.26 | 0.5 | 1.66 | 0.23 | 0.43 | 1.45 |
| Fat | 0.04 | 0.03 | 0.05 | 0.08 | 0.06 | 0.1 |
| Oxyhemoglobin | 14.06 | 12.17 | 22.44 | 3.37 | 2.92 | 5.38 |
| Deoxyhemoglobin | 73 | 34.2 | 15.48 | 59.92 | 28.11 | 12.72 |
| Oxidized Cytochrome c Oxidase | 35.64 | 14.04 | 16.2 | 2.34 | 0.9 | 1.1 |
| Reduced Cytochrome c Oxidase | 4.25 | 1.97 | 1.26 | 0.13 | 0.06 | 0.04 |

The whole tissue absorption coefficient is given in table 6.

**Table 6.**
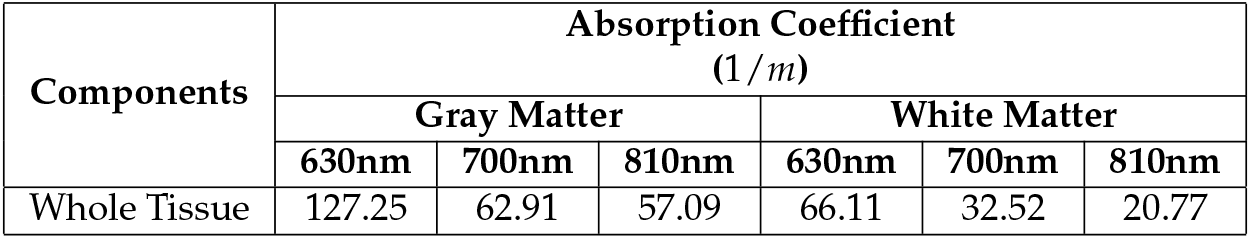
Absorption Coefficient of gray and white matter due to total contribution of all components of interest

| Components | Absorption Coefficient<br>(1/m) |  |  |  |  |  |
| --- | --- | --- | --- | --- | --- | --- |
|  | Gray Matter |  |  | White Matter |  |  |
|  | 630nm | 700nm | 810nm | 630nm | 700nm | 810nm |
| Whole Tissue | 127.25 | 62.91 | 57.09 | 66.11 | 32.52 | 20.77 |

### 2.7. Finite Element Analysis

For the simulation of the RTE (eqn.1) coupled with the bio-heat transfer equation (eqn. 8), the Finite Element Analysis used the Partial Differential Equation(PDE) toolbox of COMSOL to solve the equations based on discretization (Supplementary figure 1). In this case, the head model with the four layers has been discretized into more than 917075 tetrahedral elements forming a complete mesh. The computation of the partial differential equations is performed at each discrete unit, more precisely, at each node of the tetrahedral element of the mesh. The approximation of the solution on the entire three-dimensional head model is performed by interpolation of the data in the space between the nodes by the use of quadratic Lagrangian shape function in-built in the COMSOL software. The source position Cz, according to the 10-20 EEG system, has been chosen as the stimulation site. A point source of light with power density 500*mW*/*cm*^2^ adapted from the literature[1] was assumed at the Cz position in the head model. The source was a point source of light and was placed at the scalp at the Cz position, thus being in direct contact with the skin(figure 6). The CAD model of the adult Colin27 human head model was used with each domain(layers) assigned the optical and bioheat parameters given in tables 2 and 3.

**Figure 6.**
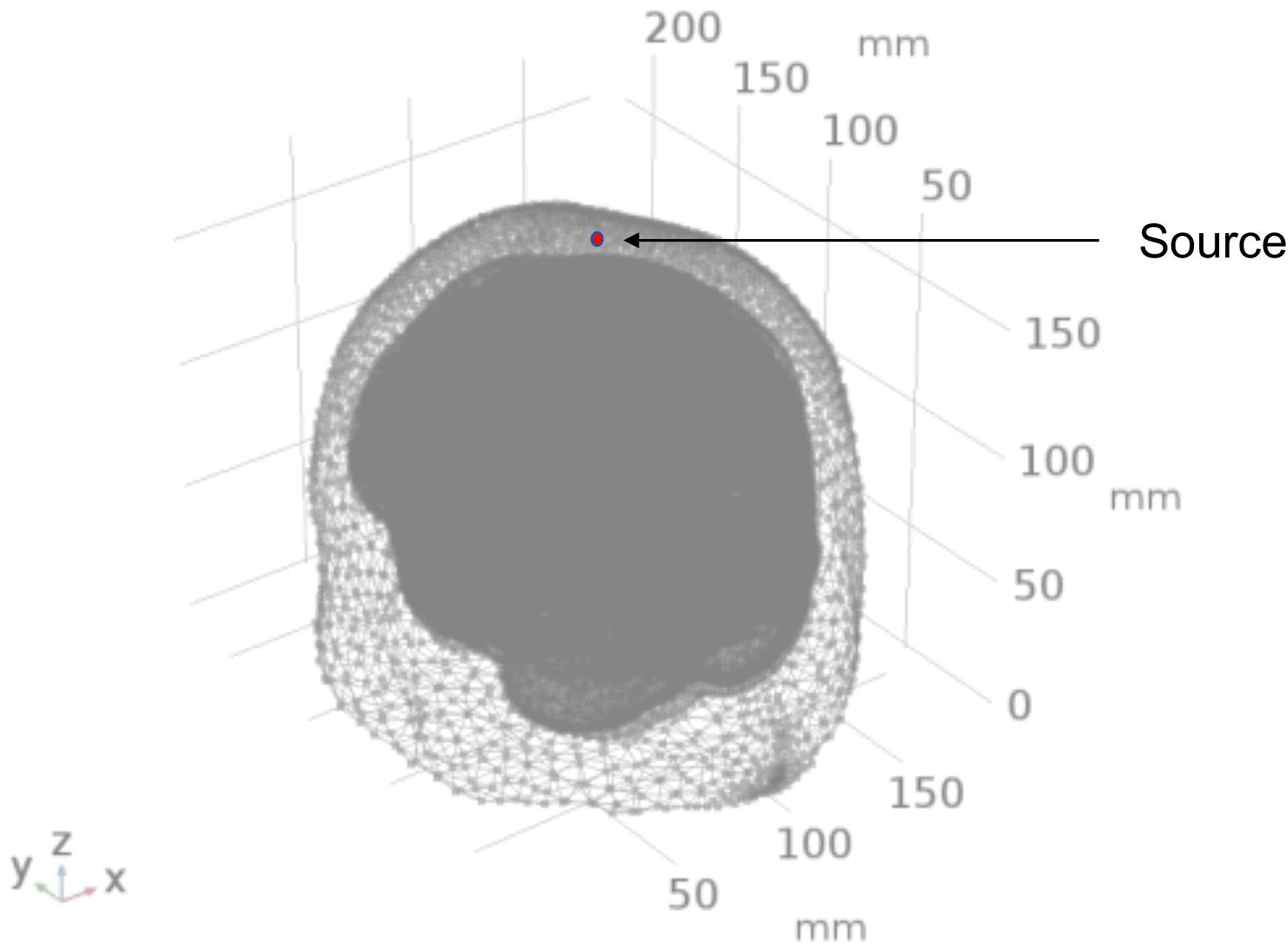
Colin27 head model with point source at Cz at scalp surface

For simulating the light interaction due to individual chromophores, we investigated the absorption in the brain tissues due to specific chromophores (based on its absorption coefficient in the brain tissue). Since scattering is a property attributed by the geometry of the medium (i.e., the brain tissue), we have assumed the same reduced scattering coefficient of the tissue during all the chromophore-specific simulation. We used both absorption and scattering properties of the skull and scalp and the CSF since we wanted to study the fluence rate at the brain tissues after the light traveled through the scalp, skull, and CSF. Thus, we initially simulated for the brain tissues’ contribution to light absorption by taking the lumped or total (due to all chromophores) absorption coefficients of the gray and white matter. The results for this specific simulation is presented as ‘Whole Tissue’. Note that the attenuation coefficient is the sum of the absorption coefficient and the scattering coefficient. Here, scattered light fluence rate is expected to be a constant (for all chromophores in the tissue) while the fluence attenuation is due to all the chromophores (absorption coefficients lumped in the tissue). To determine the attenuation due to individual chromophores of interest, we kept the tissue scattering the same (as in ‘Whole Tissue’ simulation) while determining the fluence rate attenuation due to individual chromophores in the tissue. The results were all plotted along a line cut through Cz (figure 7) crossing the layers, thereby, depicting the straight path of light into the tissue.

**Figure 7.**
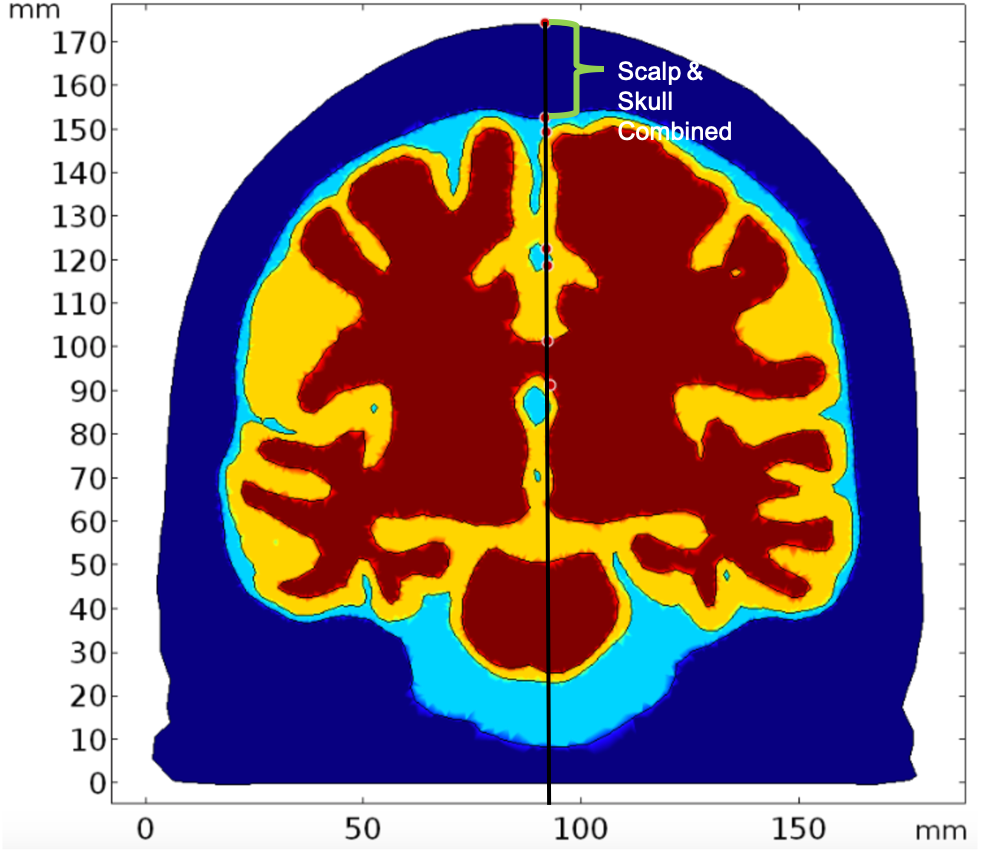
The line cut through Cz along which we analyzed the path of light(Deep blue: Scalp and skull domain, Sky blue: CSF domain, Yellow: Gray Matter domain, Red: White Matter domain)

The cutline has been drawn through the source which has coordinates 92mm,104mm, and 174mm. The x-axis on the graphs show the values of z-coordinates of the points along the cutline. Thus, values of z-coordinates decrease as the light travels further from the source placed at the scalp surface(the grid of coordinates shown in figure 6 where the z-axis is the depth).

## 3. Results

### 3.1. Photothermal effects

The optical fluence rate due to absorption and scattering by each layer is obtained from the solution of the RTE equation(7).

Figure 8 shows the normalized (natural logarithm) fluence rate for the ‘Whole Tissue’ in the tissue layers for the three wavelengths used for the study along the straight line taken through Cz. It is seen, that for wavelength 700nm and 810nm, fluence rate is comparatively higher at greater depths (less attenuation) when compared to that at 630nm, i.e., a higher penetration depth near the NIR optical window. The fractional fluence rate from the scalp surface to gray matter is shown in figure 9 where we see that minimal amount of light penetrates from the scalp through the skull and cerebrospinal fluid to the gray matter across a distance of more than 20mm along the cutline(figure 7) showing that minimal amount-around 0.2% NIR light is able to penetrate the skull.

**Figure 8.**
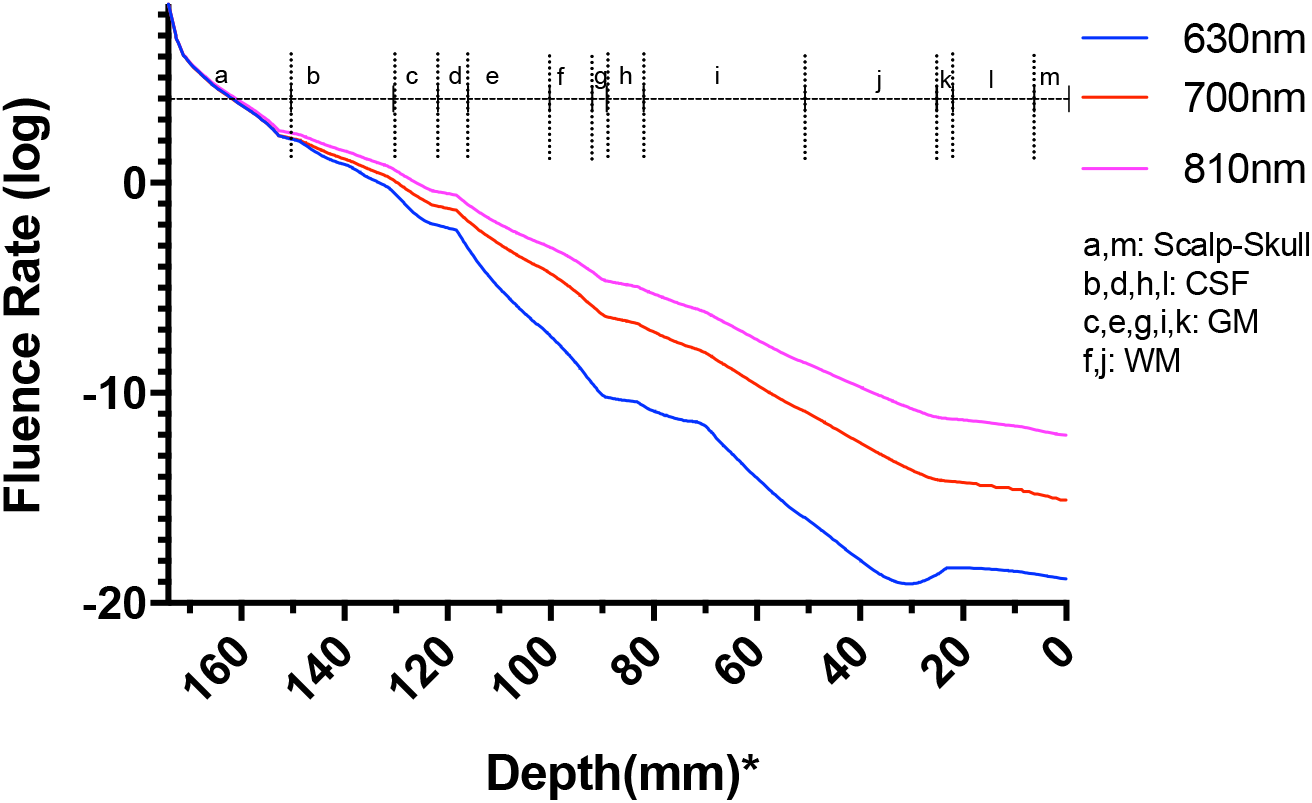
Fluence rate at the layers in the Colin27 head model for the three wavelengths; *: The numbers on the x-axis show the z-coordinates of the points on the cutline

**Figure 9.**
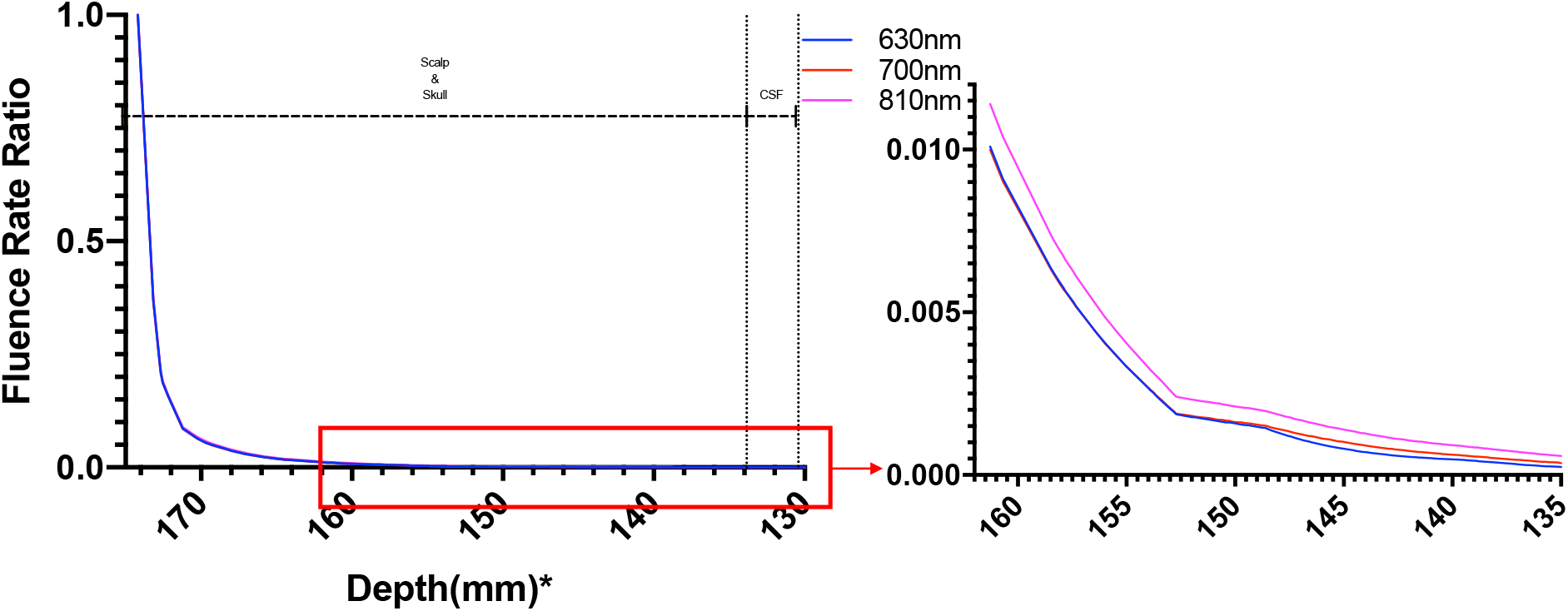
Fluence Rate Ratio from Source Gray Matter; *: The numbers on the x-axis show the z-coordinates of the points on the cutline

Figure 10 shows the power absorbed per unit volume by gray matter. The absorbed power has been assumed as the heat source for the gray matter, causing the temperature alteration.

**Figure 10.**
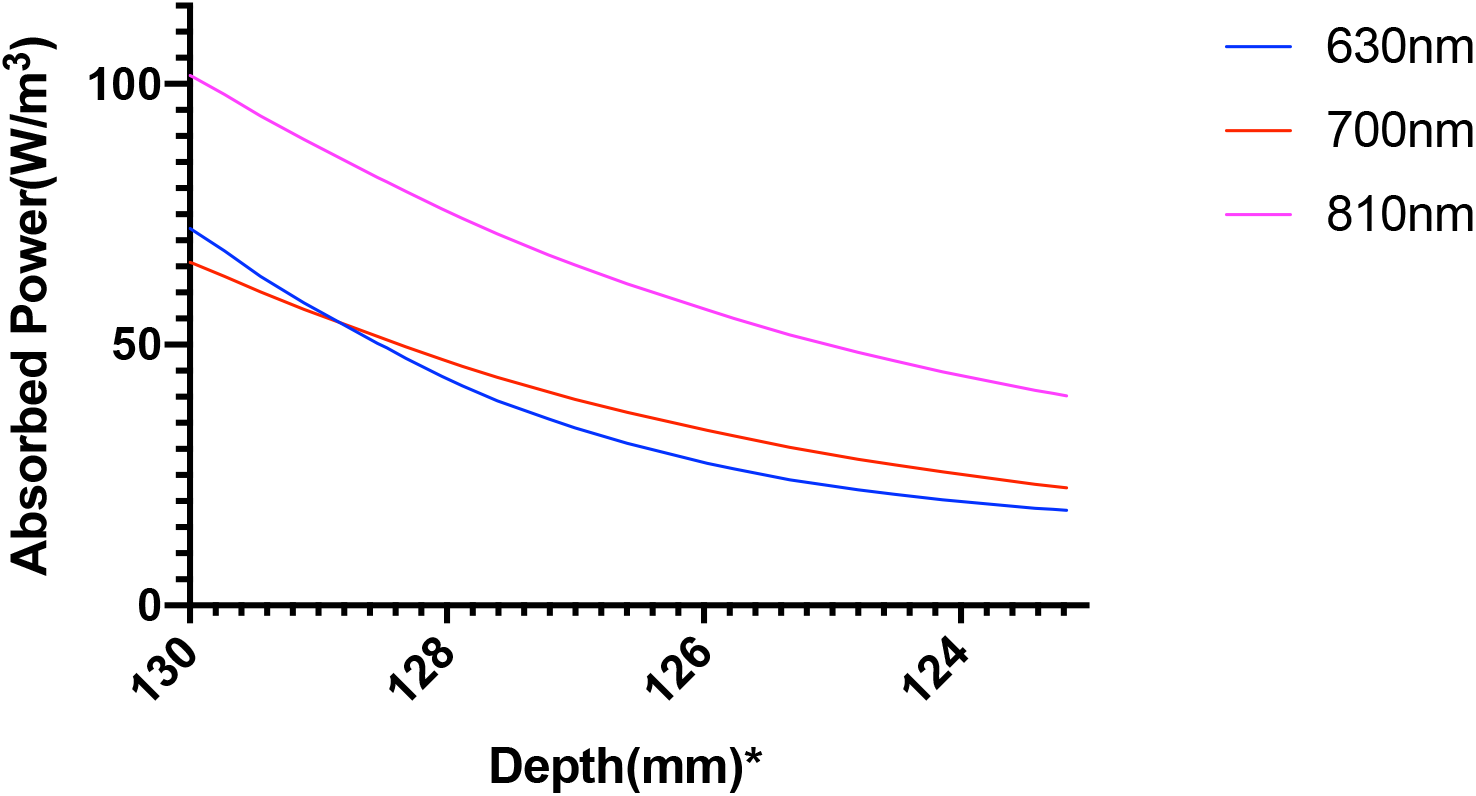
Power absorbed per unit volume(heat source) in the gray matter; *: The numbers on the x-axis show the z-coordinates of the points on the cutline

It was seen that 810nm comparatively shows a higher absorption of power at the gray matter, and thus we hypothesized that this wavelength a better choice for photothermal neuromodulation. We performed the bioheat simulation for all three wavelengths to verify our hypothesis.

The temperature along the line at the Cz location (10-20 EEG system) at different domains of the head model due to the 630nm, 700nm and 810nm optical stimulation was obtained from bioheat transfer solution, as shown in Figure 11.

**Figure 11.**
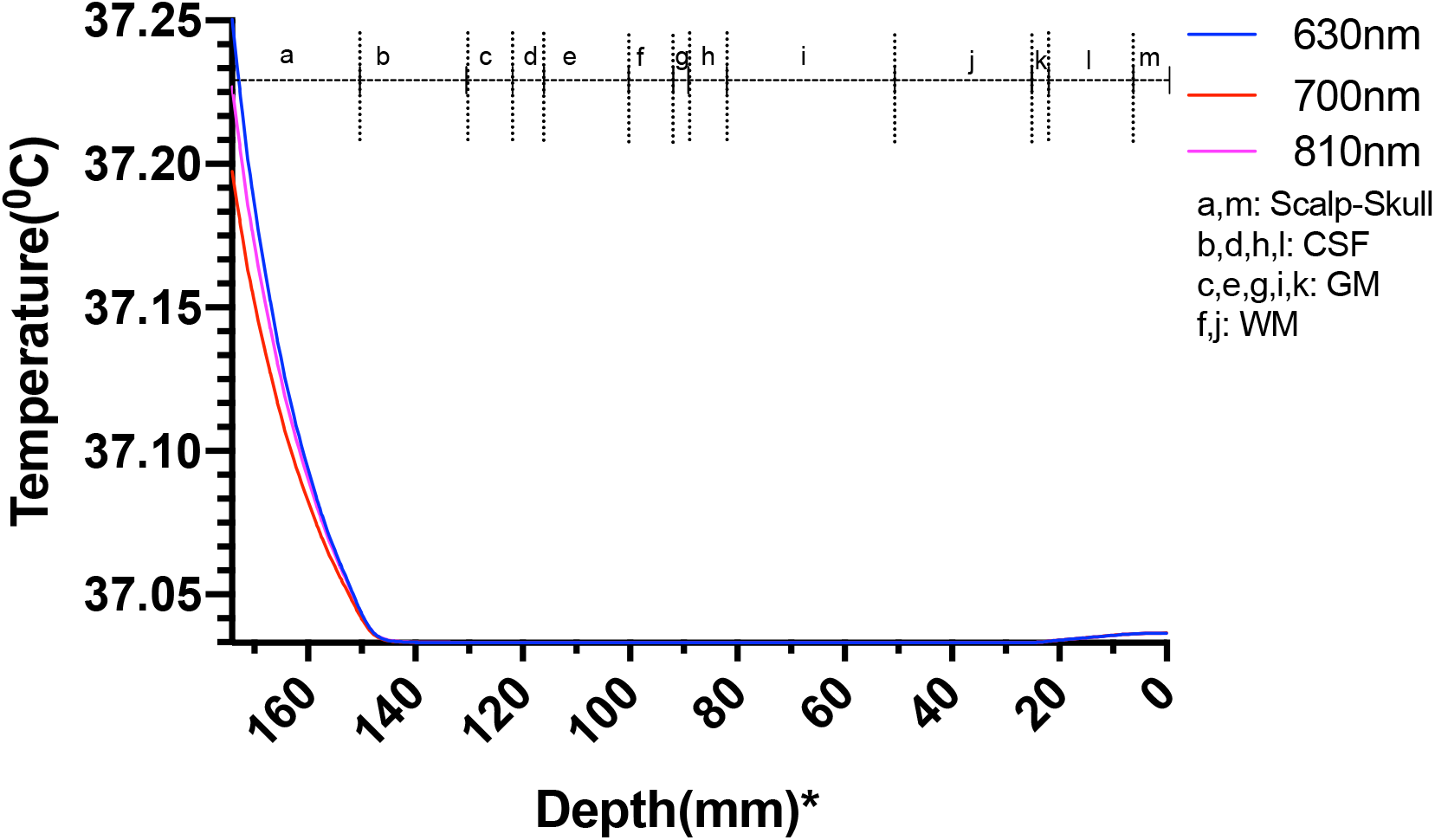
Temperature variation along the different layers in the head model plotted along the line thorugh Cz; *: The numbers on the x-axis show the z-coordinates of the points on the cutline

The results showed a temperature rise, at the scalp surface as well as at the other layers, from the average body temperature of 37 °C. The increase of temperature at the scalp at Cz is less than 0.25 °C so well within the safety limit(figure 11), but the rise of temperature at the gray matter underlying Cz area was much lower less than 0.04 °C (figure 12).

**Figure 12.**
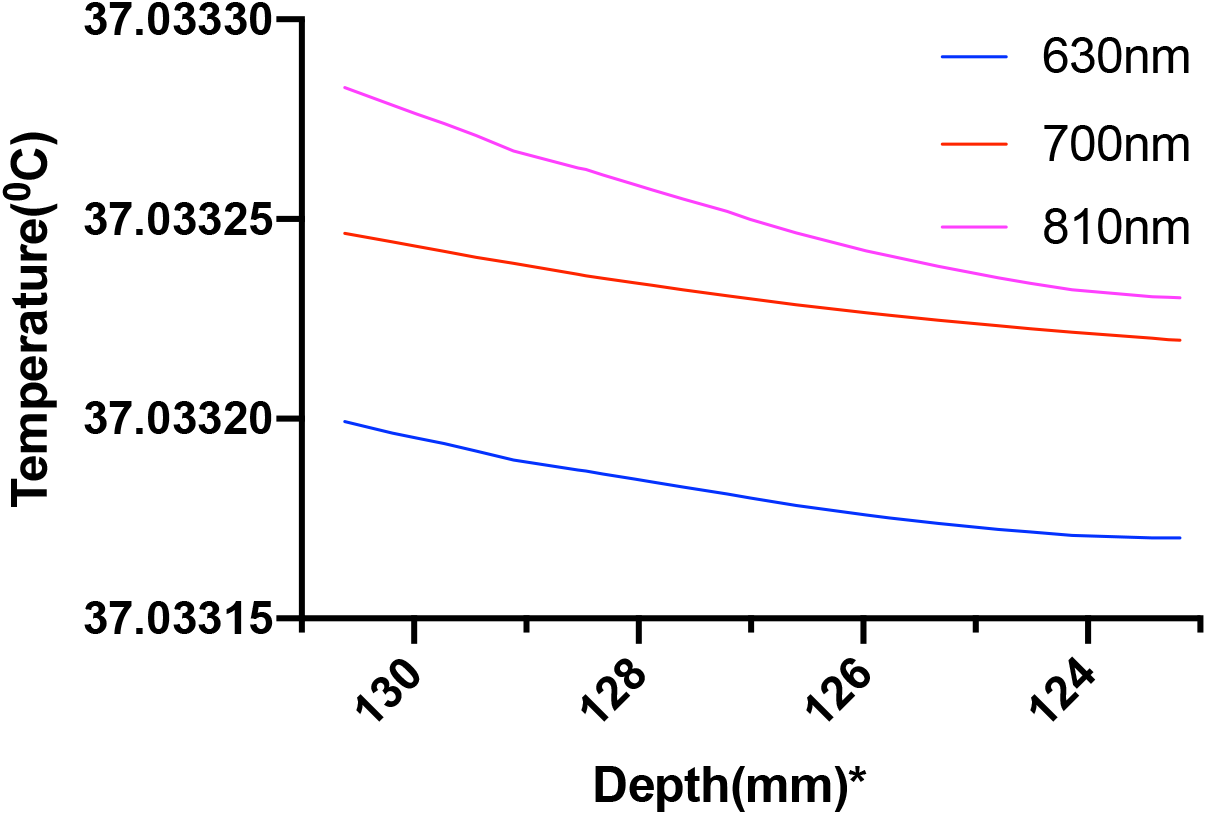
Temperature variation in the gray matter plotted along the line through Cz; *: The numbers on the x-axis show the z-coordinates of the points on the cutline

The temperature plotted over the volume of gray and white matter and represented through color map further elucidates the temperature distribution over the entire volume of the two domains, as shown in figure 13.

**Figure 13.**
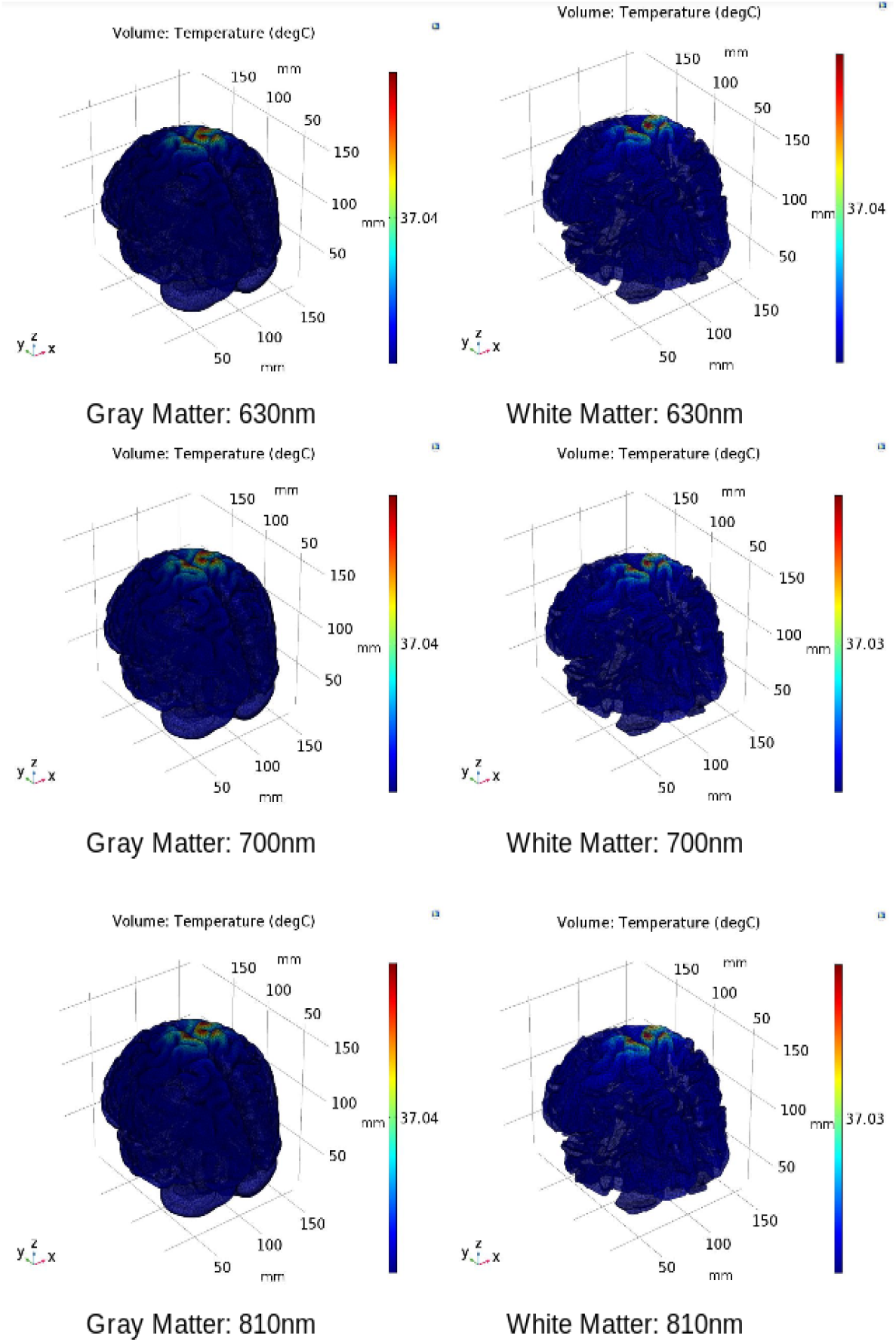
Temperature distribution in the gray and white matter volume

In figure 13, it can be seen that at all the wavelengths, there is no considerable increase in temperature in the gray and white matter and temperature is very close to the average body temperature. In both cases, the photothermal effect leading to changes in neural excitability is not expected at such a small change in temperature. Hence, we investigated the other aspect of light interaction with the neural tissue, i.e., photobiomodulation.

### 3.2. Photobiomodulation

The light interaction with the chromophores in the gray and white matter (see Table 5) was performed to analyze how the three wavelengths (630nm, 700nm, and 810 nm) in different spectral regions are absorbed in the brain tissues that can lead to photobiomodulation. The fluence rate has been computed along the cutline shown in figure 7.

The fluence rate distribution due to absorption by specific chromophores (at the gray and white matter) through the different layers of the head model is shown in figures 14, 15, 16,17, 18 and 19. Figures 14, 16 and 18. These show how the fluence rate varied in the layers along the surface normal through Cz. Our simulations support the prior findings that around 1-5% of light in the NIR spectral region reaches the gray matter [1]. We plotted the curve by taking the logarithm of the computed absolute fluence rate across different layers to better visualize the attenuation for each chromophore. An important finding when comparing between 630nm, 700nm, and 810nm is that cytochrome c oxidase contributes significantly in the attenuation of light at the gray matter, thus showing lower flux. Moreover, 810nm was found to have better depth penetration (figure 9), and so this wavelength is seen to be more promising than 630nm and 700nm for non-invasive brain stimulation.

**Figure 14.**
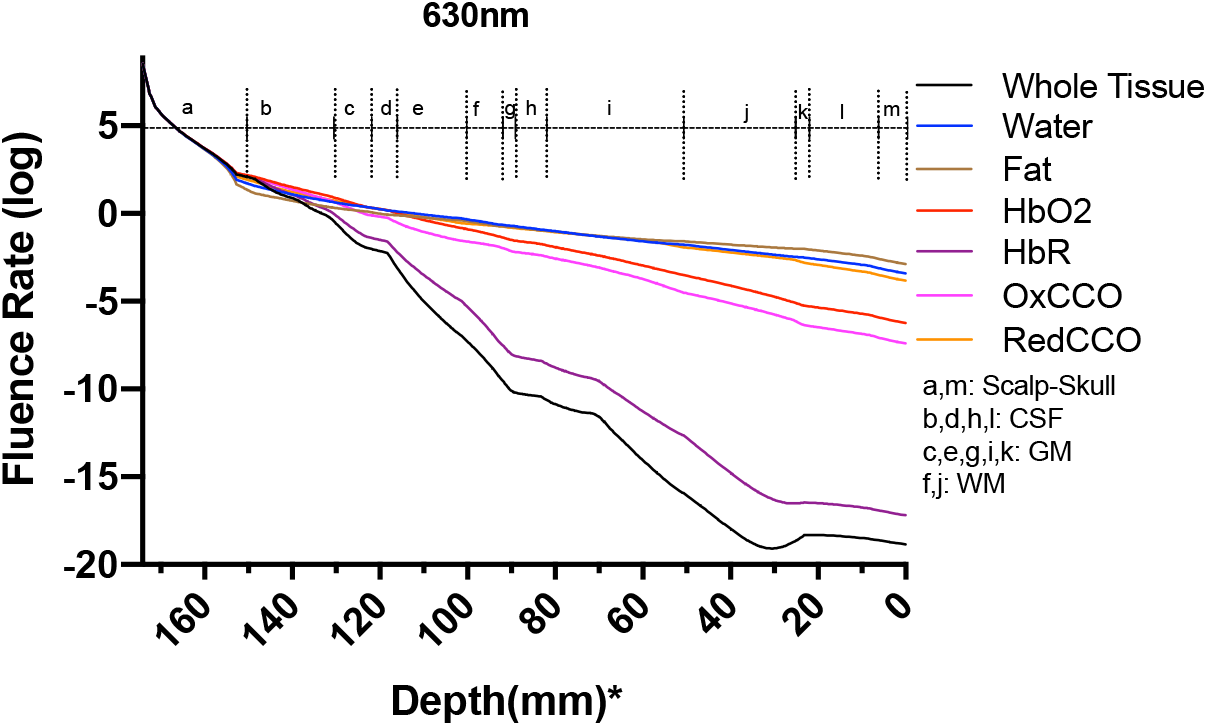
Fluence rate through the different layers of the head model at 630nm due to individual components at the gray and white matter; *: The numbers on the x-axis show the z-coordinates of the points on the cutline

**Figure 15.**
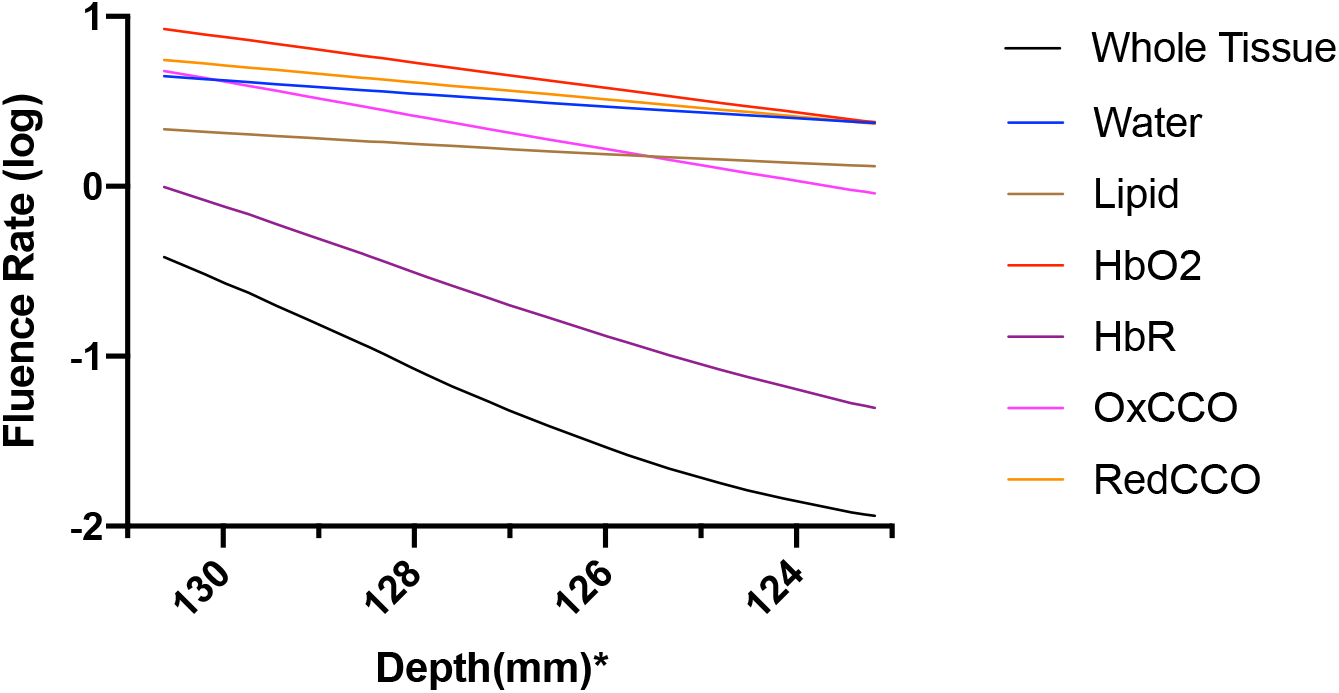
Fluence rate at 630nm in the gray matter due to individual components; *: The numbers on the x-axis show the z-coordinates of the points on the cutline

**Figure 16.**
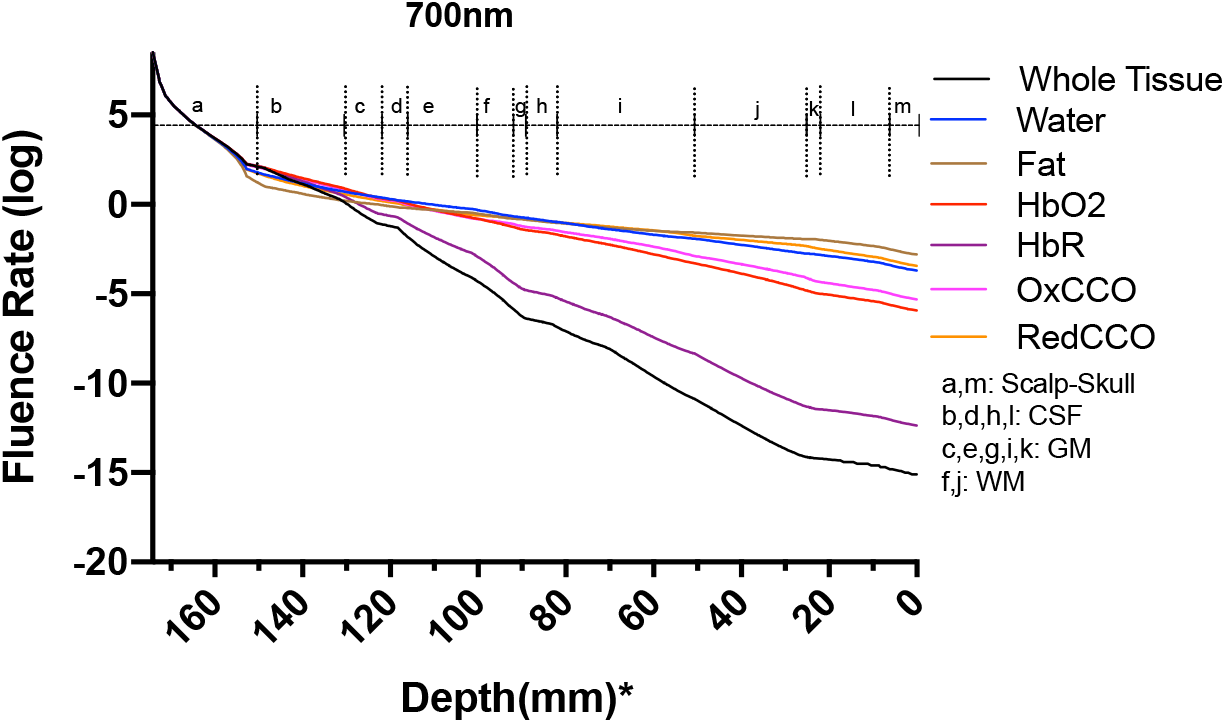
Fluence rate through the different layers of the head model at 700nm due to individual components at the gray and white matter; *: The numbers on the x-axis show the z-coordinates of the points on the cutline

**Figure 17.**
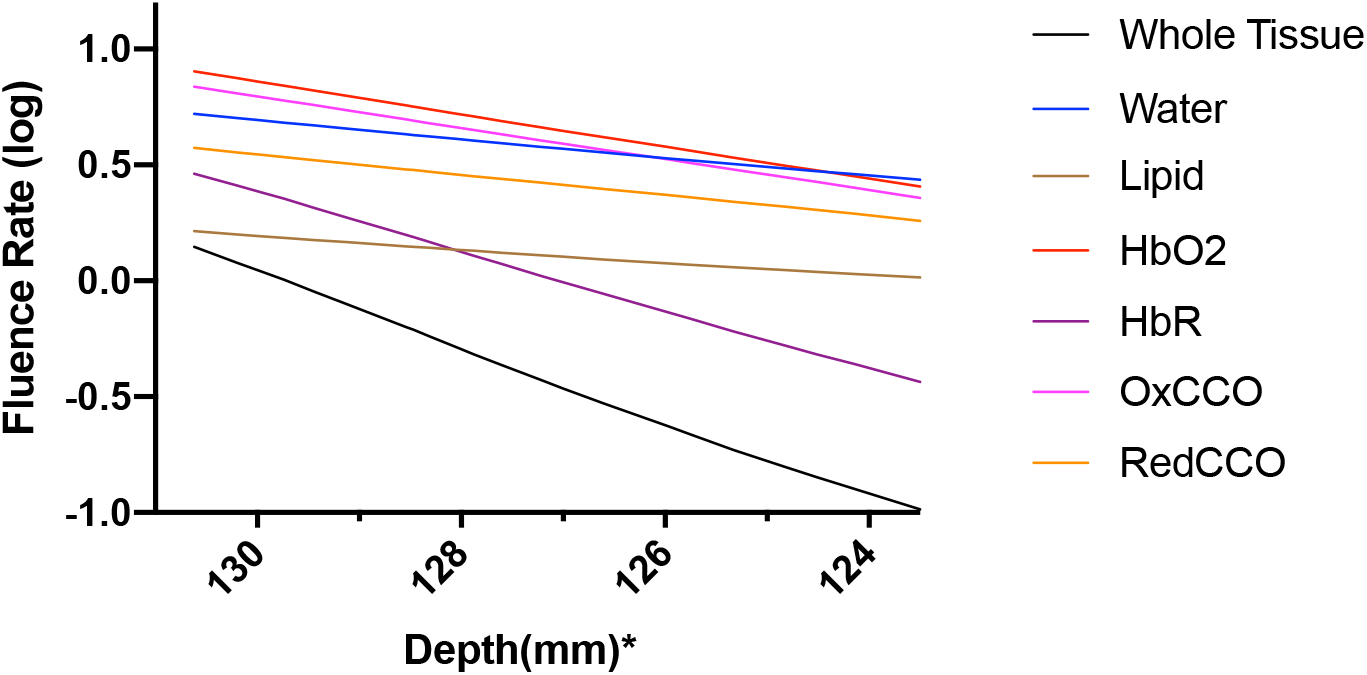
Fluence rate at 700nm in the gray matter due to individual components; *: The numbers on the x-axis show the z-coordinates of the points on the cutline

**Figure 18.**
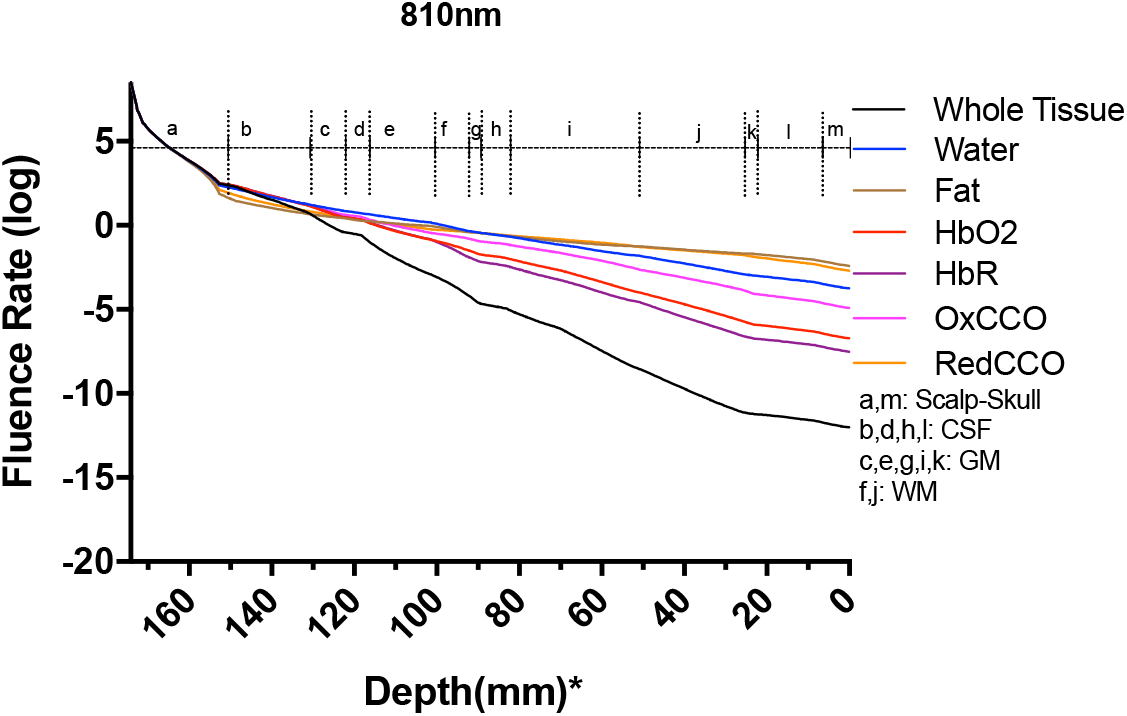
Fluence rate through the different layers of the head model at 810nm due to individual components at the gray and white matter; *: The numbers on the x-axis show the z-coordinates of the points on the cutline

**Figure 19.**
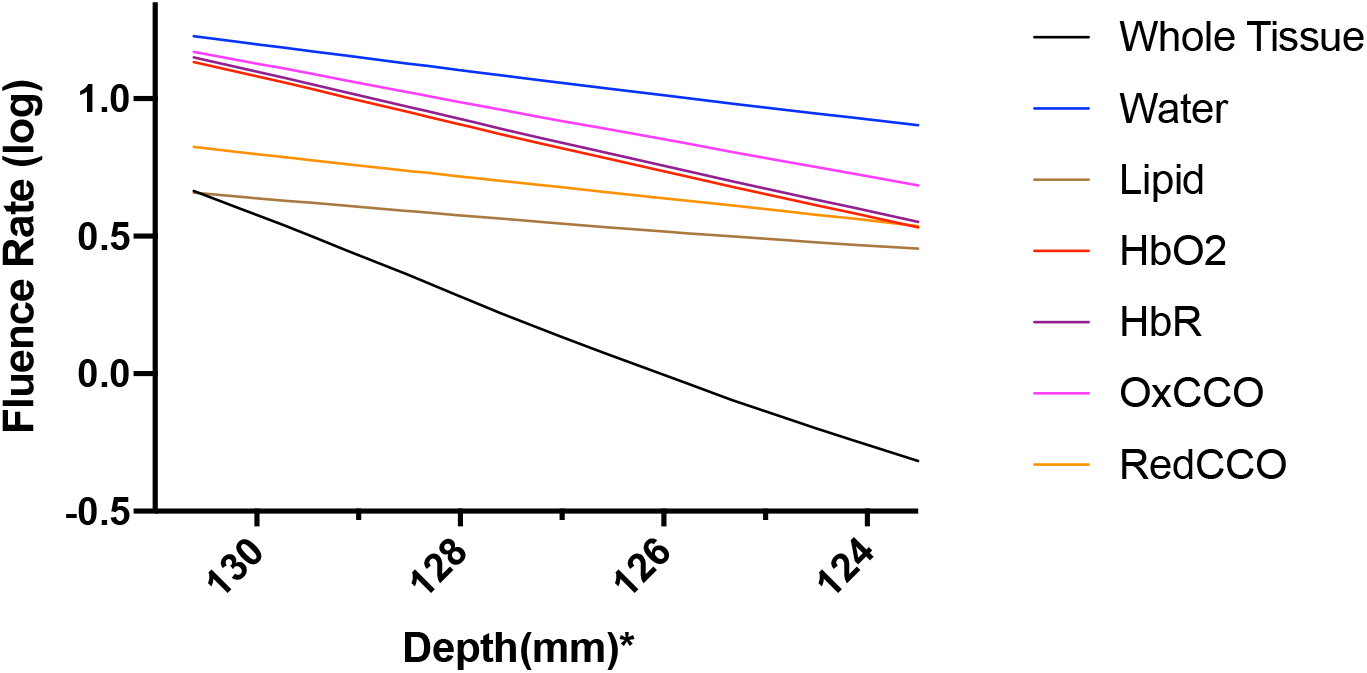
Fluence Rate at 810nm in the gray matter due to individual components; *: The numbers on the x-axis show the z-coordinates of the points on the cutline

## 4. Systematic Model Errors

The simulated model shown here is a realistic head model based on the Colin27 head atlas. The mathematical model could show errors, mainly due to simplifications. The errors may be as follows:

- In our model, the brain has been assumed as a highly scattering medium which is not true for CSF which is a low scattering medium where RTE can produce erroneous results[57](Comparison between RTE and Monte Carlo has been shown in Supplementary material).
- Another error is related to computational limitation during the FEM modeling and discretization.
- There were limitations of accessible memory. Although enhancing resolution leads to better convergence of FEM results[58], but computational limitation restricted us to the standard COMSOL mesh refining process.
- The reflection effects due to light interaction have been excluded from the simulations to focus on light interaction with tissues due to absorption and scattering.
- The optical properties of the tissues in the head model vary significantly based on prior works. We selected a set of optical parameters from review literature[39][56][49]. We did not consider chromophores in the skin such as melanin, lipofuscin.
- The simulation of the Bioheat Transfer assumed that the heat loss at the skin surface is due to convection and radiative heat loss was considered insignificant at that temperature.

## 5. Discussion and Conclusion

The computational pipeline aimed to investigate the temperature change induced by the light absorption at the three wavelengths, 630nm, 700nm, and 810nm, as well as the absorption by the chromophores in the neural tissue. In this multiphysics modeling of light diffusion with bioheat transfer, we found that the temperature change in the scalp is well within 1 degree Celsius as reported by Chaieb and colleagues [1] for a light source of power density 500*mW*/*cm*^2^ at the scalp surface. As the light gets attenuated while propagating through the skull and cerebrospinal fluid (CSF) to reach the gray matter(0.2% reaches the gray matter; hence less than 1%), the low fluence rate leads to insignificant heating in the gray matter. Here, we assumed the initial body temperature at 37°C, and the temperature increase at the Cz area was found to be around 0.033°C at the gray matter-CSF interface. The sharp decrease in the fluence rate as the light propagated further into the gray matter is shown in figure 12.

Prior works on brain temperature that was assessed on resting clinical patients showed an average brain temperature ranging around 36.9*±* 0.4°C[59]. Brain activity has been shown to be associated with the rise in brain temperature. Studies have suggested that temperature changes of even less than 1°C can result in functional alterations in the various areas of the nervous system[60], indicating the high thermal sensitivity of the brain. Thus, the temperature is an important active and dynamic variable that can modulate brain activity and needs to be monitored during stimulation. The brain, being at a typically higher temperature than the body, is cooled down by perfusing blood, which was considered in our modeling of thermal changes through bioheat transfer. Here, blood perfusion acts as a heat sink, thus cooling the brain down. Therefore, the temperature change in biological tissue is significantly dependent on the bioheat, which is further dependent on the heat source and the blood perfusion sink. The simulated results showed insignificant temperature change (0.033°C) to cause photothermal neuromodulation. Hence, the chromophore simulations suggest a possible photobiomodulation effect of the NIR light interaction with the tissue. The results obtained from the simulation of the absorption by each chromophore, including lipid and water, elucidated the fact that besides water and lipid, light attenuation in the gray matter is due to the absorption of NIR light by the reduced and oxidized CCO. In fact, for 630nm, 700nm, and 810nm wavelengths, we found that the two forms of CCO are the two major contributors to light attenuation besides water, lipid, and hemoglobin. Thus, we can conclude from the three categories of data that neuromodulation of the gray matter by photothermal effect is not significant with 500mW cm^−2^ at the scalp surface at 630nm and 700nm (red spectral region) and 810nm (near-infrared spectral region). However, the biochemical effects of CCO absorption need further investigation in conjunction with the heating effects since a small, steady state temperature change can affect the kinetics of photobiomodulation. Our simulation data comparing the fluence rate attenuation among 630nm, 700nm, and 810nm also showed that 810nm has higher penetration depth than the 630nm and 700nm, which supports the use of tNIRS for non-invasive brain stimulation.

## Acknowledgments

We thank the reviewers for their insightful comments and suggestions that improved the work presented in the paper. We also thank Steffy Rodriguez (undergraduate student of Biomedical Engineering at the University at Buffalo) for proofreading the manuscript.

## Abbreviations

The following abbreviations are used in this manuscript:

PBM: Photobiomodulation
CCO: Cytochrome c oxidase
ROS: Reactive Oxygen Species
FEM: Finite Element Method
NIR: Near Infrared
RTE: Radiative Transfer Equation
PDE: Partial Differential Equation
tNIRS: transcranial Near Infrared Stimulation

